# Frequency and Characterization of Rare Histologic Subtypes in Canine Invasive Urothelial Carcinoma

**DOI:** 10.1101/2025.05.13.651218

**Authors:** Megan P. Corbett, Mohamed Elbadawy, Andrew P. Woodward, John C. Cheville, Hannah Wickham, Hannah Nicholson, Michael Catucci, Bryan J. Melvin, Basant Ahmed, Christopher Zdyrski, Lilian J. Oliveira, Elizabeth W. Howerth, Aleksandra Pawlak, Olufemi Fasina, Jonathan P. Mochel, Karin Allenspach

## Abstract

The histologic and molecular heterogeneity of human muscle-invasive bladder cancer (MIBC) is a major contributor to poor treatment outcomes. While most cases of MIBC are diagnosed as conventional urothelial carcinoma (UC), there is growing recognition of histologic subtypes and divergent differentiation within conventional UC. This is clinically significant, as some require modification of therapy and some are associated with more aggressive behavior. Spontaneously occurring UC in dogs has been shown to exhibit histologic and molecular features that closely resemble those of human MIBC. In this study, we evaluated 31 canine UC tumor samples for histologic subtypes and divergent differentiation. Slides were reviewed by a human uropathologist and three board-certified veterinary pathologists and assessed for expression of uroplakin III and E-cadherin. All tumors were classified as high-grade UC. Fifteen cases were identified as conventional UC. Among the remainder, eight displayed glandular differentiation, four were classified as sarcomatoid UC, two showed squamous differentiation, and one case each was classified as large nested and tubular and microcystic subtypes.

In summary, this study found a higher frequency of certain histologic subtypes and divergent differentiation in canine UC—particularly sarcomatoid UC and UC with glandular differentiation—compared to previous reports in both canine UC and human MIBC.

**Conclusion:** The relatively high prevalence of the sarcomatoid UC subtype in dogs observed in this study suggests that canine UC may serve as a valuable translational model for evaluating novel therapeutic agents, particularly for this rare and aggressive variant in humans.

---

The most common type of urinary bladder cancer in humans is urothelial carcinoma (UC), which is currently one of the top ten most common solid organ malignancies worldwide, with approximately 600,000 new cases and 220,000 deaths in 2022 ^8^. The disease is heterogeneous, with non-muscle-invasive bladder cancer generally having better outcomes, while aggressive muscle-invasive bladder cancer (MIBC) is associated with a poorer prognosis ^47^. The significant intra-tumor heterogeneity described in human bladder cancer has prompted the development of various classification systems based on molecular subtyping, such as the Lund Taxonomy System ^20^, the World Health Organization (WHO)-^41^, TCGA-^62^, MD Anderson-classification system ^22^ and the consensus statement of the European Society for Urology ^2,23^. Overall, all of these molecular classification systems converge on the principal criteria of tumors that express either predominantly basal markers (such as CD44, CK5/6, and CK14), indicating their cellular origin from the basal and intermediate layers of the urothelium, or luminal markers (such as FOXA1, GATA3, UPK and CK20), indicating their cellular origin from urothelial umbrella cells ^50^. These classifications have been linked to pathological grade, invasiveness, as well as treatment responses, and clinical outcomes ^49^, with an overall agreement between a higher grade and basal features of tumors being associated with a more progressive disease course. Critically, however, molecular subtyping does not generally determine histological subtype or divergent differentiation in MIBC, which are features that are still most commonly identified by experienced pathologists upon histological review ^7,36,64^. Additionally, molecular subtyping is not used clinically whereas reporting subtypes and divergent differentiation is the standard of care in human bladder cancer. Histological subtypes of UC in this context are defined as tumors that develop within the bladder that also contain some components or be entirely composed of a morphologically distinct carcinoma from conventional UC such as neuroendocrine carcinoma or sarcomatoid carcinoma. Furthermore, divergent differentiation (also referred to as aberrant differentiation) is used when areas of the urothelial tumor exhibit a different histogenesis such as squamous or glandular differentiation. These histological subtypes and divergent differentiations are important to recognize and report, as some subtypes have an association with more aggressive behavior and poorer response to conventional chemotherapy, immune checkpoint inhibitors, and antibody-drug conjugates^34^. In 2022, the WHO updated its classification of urinary bladder cancers to include histological subtypes of UC (formerly called histological variants)^47^. The poor treatment outcome in some histological subtypes of UC has been attributed to the lack of models that faithfully represent the phenotypic and molecular heterogeneity within MIBC, making it difficult to translate pre-clinical efficacy data of drugs to human patients ^29,73^. For instance, the use of murine bladder cancer models to identify effective treatment for these tumors has been limited because they do not accurately represent the biological behavior of human MIBC, including important features such as metastatic tendencies ^51^. Additionally, immunotherapeutic approaches cannot be easily studied in xenotransplant mouse models that lack an intact immune system, which is imperative for pre-clinical efficacy evaluation of immune checkpoint inhibitor and antibody-drug conjugate therapies. On the contrary, dogs with naturally occurring UC have shown similar heterogeneity, molecular subtype, clinical, and metastatic behavior, as well as a comparable tumor microenvironment to those in humans with MIBC ^16,57,69^. Interestingly, it has recently been established that canine UC can present with histological subtypes similar to those in human medicine, including sarcomatoid, plasmacytoid, rhabdoid, and neuroendocrine variants ^14,25,33^. In addition, these histological subtypes could be associated with similarly aggressive behavior as in humans, warranting further characterization of histologic subtypes and divergent differentiation of UC in dogs ^14^.

In the current study we aimed to characterize the histological subtypes and divergent differentiation of canine UC in a series of 31 cases. To adhere to the newly devised human classification systems ^7^, a board-certified medical uropathologist and three board-certified veterinary pathologists collaborated to review the canine tumor histology. Immunohistochemical staining was used to demonstrate non-UC morphology within a subset of the tumors. We found that histological subtypes and divergent differentiation of canine UC can be readily identified and may be present at a higher prevalence than previously reported.

## MATERIALS and METHODS

### Canine UC tumor histopathological assessment

Canine UC tumor samples that had been collected during diagnostic cystoscopy or necropsy from 31 dogs at three veterinary teaching hospitals were evaluated. Samples from Iowa State University (14/31) and the University of Georgia (14/31) were retrieved from archival material owned by the respective institutions, and owner consent had been obtained at time of treatment. Samples from Purdue University (3/31) were collected in conjunction with a prospective study (IACUC protocol 1111000169) with owner consent. One dog had been sampled at two different time points (first sample at 11-years-old and second sample at 12-years-old) and another dog had a large tumor sample split between two separate slides, resulting in 33 slides. Images of hematoxylin and eosin (H&E) stained slides were captured using an ECHO Revolution microscope (RNSD1000, Echo Apico Co.), Leica Versa 8 model DM6 B scanner, or a Leica Aperio GT 450 scanner at 40x magnification. Histopathological examination, grading, and subtyping were performed in a blinded fashion (**Supplemental Table S1**). Slides were evaluated by a board-certified medical uropathologist alongside 3 board-certified veterinary pathologists (Dr. Cheville, Mayo Clinic, Rochester, MN, Dr. Oliveira, College of Veterinary Medicine, University of Georgia, Dr. Corbett, College of Veterinary Medicine, University of Georgia, and Dr. Fasina, College of Veterinary Medicine, Iowa State University). Signalment and clinical data for canine cases are summarized in Supplemental Table S1. The presence of apparent metastasis was recorded from the electronic medical records if metastasis was definitively identified by necropsy, radiographs, or computed tomography. Furthermore, the Mayo Clinic Cystectomy Registry in Rochester, MN was assessed to provide the frequency of bladder cancer subtypes in humans. The registry follows over 3500 patients that have undergone a cystectomy for bladder cancer at the Mayo Clinic since 1980. The number and types of histological subtypes and divergent differentiations identified in this canine case series were subsequently compared with data previously reported in the human and veterinary literature, as well as with the incidence found in the Mayo Clinic Cystectomy Registry (**Table 1**).

**Table 1.**
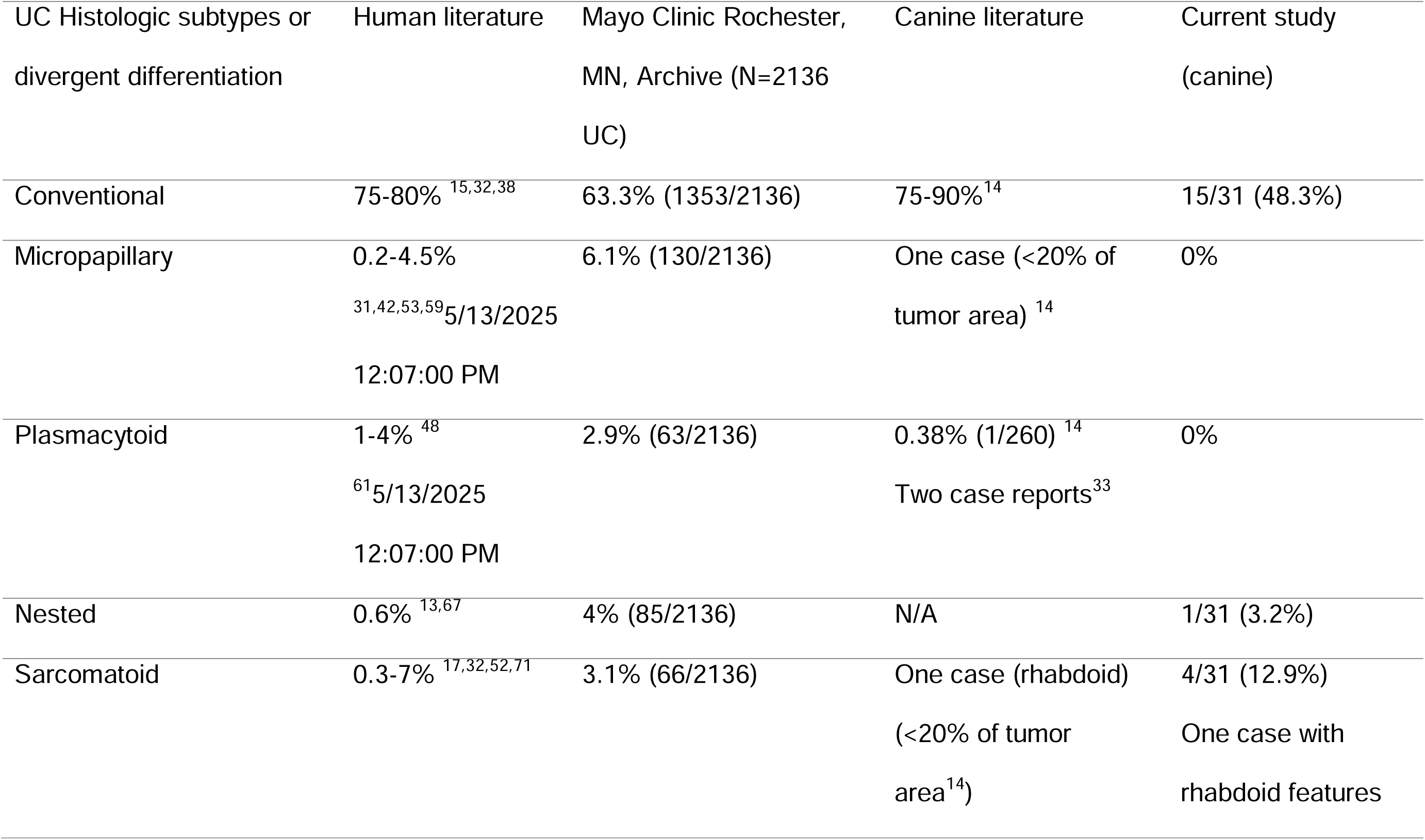
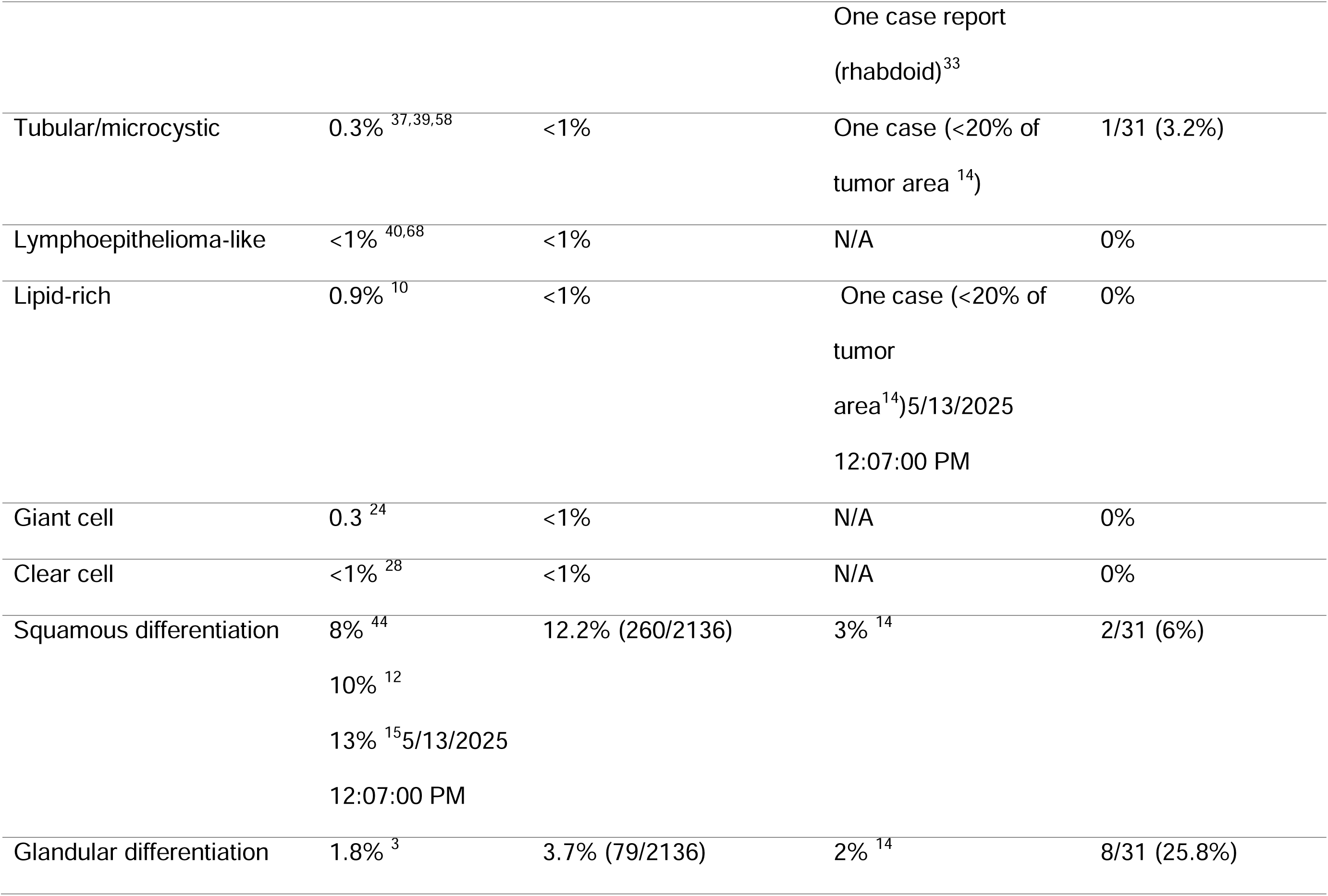
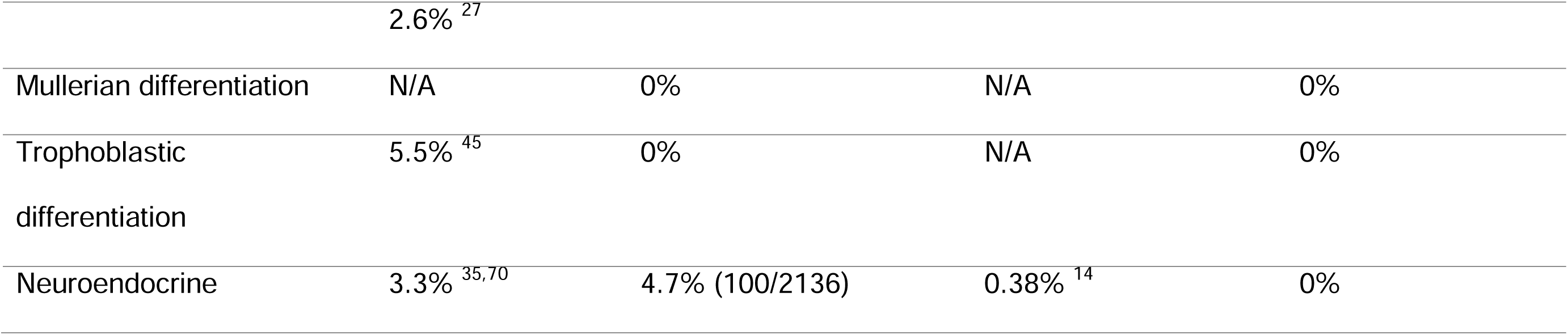
Incidences of histologic subtypes and divergent differentiation of human MIBC and canine UC according to the WHO/ISUP classification system of human bladder tumors^71^.

### Immunohistochemistry

Representative sections of each canine UC tumor subtype and divergent differentiation were further characterized by immunohistochemistry (IHC), as described in Supplemental Table S2. Due to block limitations, the following numbers of cases were used to further characterize each subtype/divergent differentiation using IHC: Six/15 conventional UC, 3/8 UC with glandular differentiation, 3/4 sarcomatoid UC, 1/2 UC with squamous differentiation, 1/1 large nested UC, and 1/1 microcystic/tubular UC. Formalin fixed, paraffin embedded sections were processed according to the standard operating procedures of each institution. Uroplakin III IHC was performed using a Leica Bond Max (Leica Biosystems) automated stainer at Cornell University. Vimentin (V9) and E-cadherin IHC was performed using a Biocare Intellipath (Biocare Medical) automated slide stainer at the University of Georgia. Chromogens were 3,3-diaminobenzidine (DAB) (BioCare Medical) for Uroplakin III and vimentin (V9) and Warp Red (BioCare medical) chromagen for E-cadherin, with hematoxylin counterstain for all slides. Images were obtained from digitized slides scanned using a Leica Aperio GT 450 scanner, Leica Versa 8 model DM6 B scanner, or Leica Aperio AT2 scanner at 40x magnification, captured with ImageScope software (version 12.4.3.5008, Leica Biosystems), and autocontrast and white balance correction applied to all images using Adobe Photoshop 2022 software (version 23.1.0).

### Data Analysis

Data analyses were conducted in R version 4.4.2. 2024 ^46^. Survival time was calculated from clinical records as either the amount of time elapsed from clinical diagnosis or reported onset of clinical signs to time of death or loss to follow up. To summarize the prevalence of metastasis by variant, a hierarchical logistic regression model was generated using the package ‘brms’^9^, with weak prior information. The posterior-predicted prevalence from the model, corresponding to the expected prevalence for each variant, was visualized to summarize evidence of between-variant variation. Due to the small sample size, imbalance, and opportunistic nature of the cohort, this analysis is intended as descriptive and cannot support any general claims or conclusions ^5^. Survival time from presentation, across all variants, was summarized using the Kaplan-Meier method as implemented in package ‘survival’^46^. The amount of data was not sufficient to support assessment of survivorship by variant. For simple prevalence statements at the cohort level, confidence intervals for the prevalence were obtained using the Wilson method ^66^.

### Data Availability

All data and codes from the present study are available at https://doi.org/10.5281/zenodo.15151129.

## RESULTS

The present study evaluated 31 canine UC tumor samples (14 males and 17 females, all surgically neutered except for one male and two female intact dogs) for histological subtypes and divergent differentiation. Similar histological subtypes and patterns of divergent differentiation were identified in the canine tumors as those observed in human MIBC, as summarized in **Table 1**.

Of the 31 dogs, 10 (32%) had definitively papillary tumors characterized by neoplastic urothelial cells supported by distinct fibrovascular stalks. The remaining samples were either plaque-like or too limited in size to definitively determine their architecture. Histopathological review determined that 15 of the 31 cases (48.3%) represented conventional UC without any identifiable subtype (**Table 1**; **Fig. 1a**). One dog, sampled twice a year apart, had papillary conventional UC in both samples, with no subtype identified. Microscopic features included eosinophilic cytoplasmic inclusions (Melamed-Wolinska bodies), clear cytoplasmic vacuoles with nuclear displacement, and frequent microcysts (also referred to as lumens) filled with variably eosinophilic to basophilic amorphous material (**Fig. 1a, arrowheads**).

**Figure 1.**
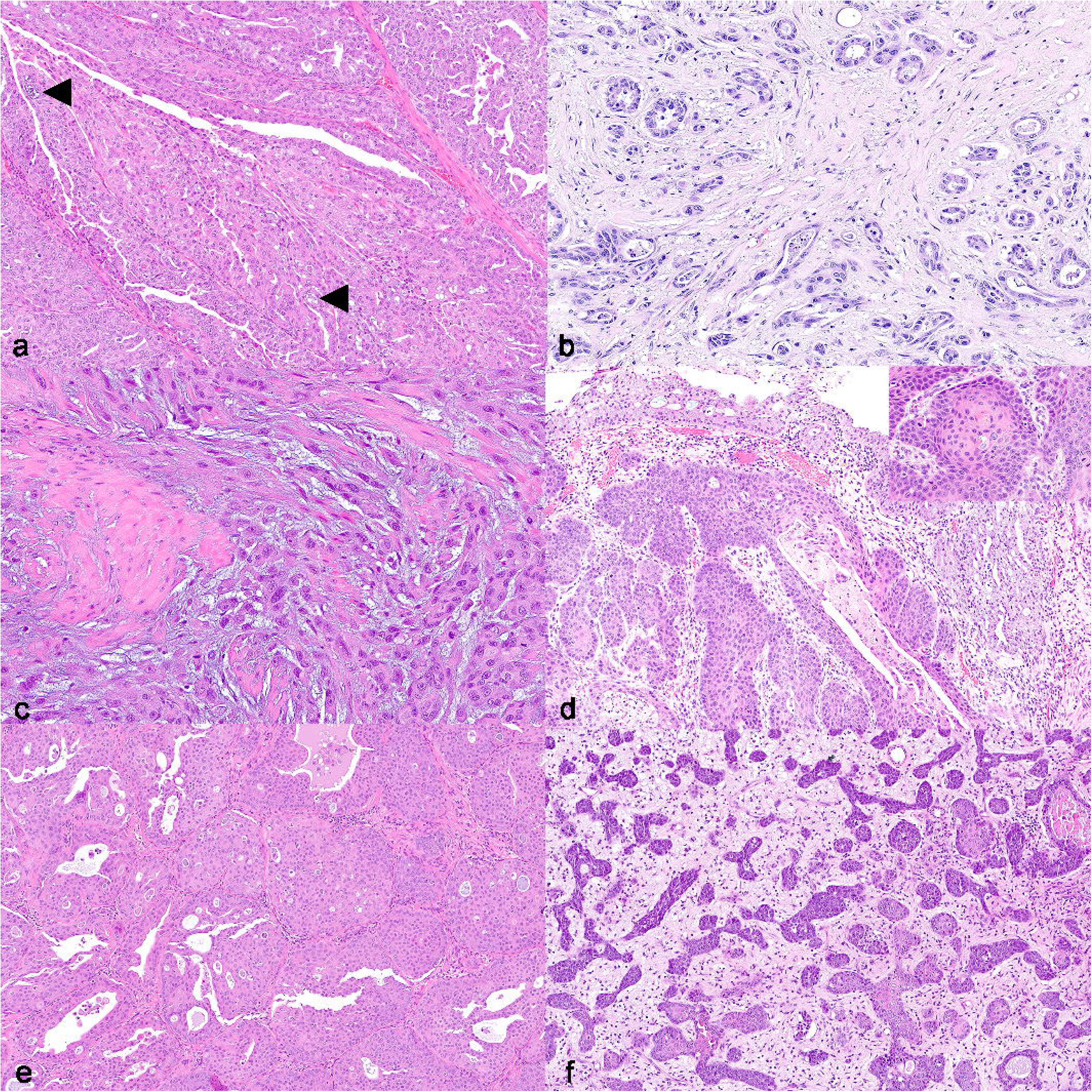
Urothelial carcinoma (UC) with varying histologic morphologies in the urogenital tract of dogs, hematoxylin and eosin (HE). a) Conventional papillary UC composed of fronds of neoplastic urothelial cells supported by a fibrovascular stalk. Scattered microcysts (lumens) filled with basophilic, mucinous material are often scattered throughout the neoplasm (arrowheads). Neoplastic urothelial cells have distinct cell borders, abundant eosinophilic cytoplasm, and round to ovoid nuclei with prominent nucleoli. Case 7. b). UC with glandular differentiation showing acinar and cribriform architecture lined by simple to pseudostratified epithelium, surrounded by abundant scirrhous stroma. Case 9. c) Sarcomatoid UC demonstrating mesenchymal differentiation with a spindled appearance, surrounded by myxoid matrix. Case 14. d) UC with squamous differentiation transitioning from adjacent conventional urothelial carcinoma in the ureter. Urothelial cells with squamous differentiation have distinct intercellular bridges (desmosomes) and hypereosinophilic, glassy cytoplasm (keratin) that form small “keratin pearls” (inset, top right). Case 28. e) High-grade, invasive, large nested subtype of UC forming large, complex nests in the superficial lamina propria. The nests are haphazardly arranged, irregular in size and shape, and form a jagged tumor-stroma interface. There is minimal stromal reaction. Case 30. f) Microcystic/tubular subtype of UC forming small, irregularly shaped nests with scattered lumens, giving a characteristic “puzzle-piece” appearance. Case 31.

Sixteen samples had histological subtypes or divergent differentiation. Some subtypes—such as sarcomatoid UC (12.9%; 95% CI [0.05, 0.29]) and UC with glandular differentiation (25.8%; 95% CI [0.14, 0.43])—were more prevalent in this cohort than reported in prior human and canine studies (**Table 1**). In one case, conventional UC was found in one slide, while UC with glandular differentiation was present in another slide from the same tumor, highlighting the potential impact of tumor heterogeneity on subtype detection.

UC with glandular differentiation was characterized by the formation of tubule- and gland-like structures lined by one to three layers of pseudostratified neoplastic urothelial cells, resembling intestinal adenocarcinoma (**Fig. 1b**). Sarcomatoid UC (**Fig. 1c**) was identified in 4 of 31 samples (12.9%) and featured extensive infiltrative regions of urothelial cells with wispy cytoplasm, indistinct borders, and elongated nuclei often surrounded by basophilic (mucoid) matrix. One tumor showed rhabdoid morphology, with nuclear rowing and abundant, eosinophilic, strap-like cytoplasm.

Two of 31 tumors (6%; 95% CI [0.02, 0.21]) had squamous differentiation (**Fig. 1d**), marked by areas of neoplastic urothelium transitioning into islands of epithelial cells with well-defined junctions, hypereosinophilic, glassy cytoplasm (keratinization), keratohyalin granules, and keratin accumulation (**Fig. 1d**, inset). A large nested subtype was observed in 1 of 31 tumors (3.2%; 95% CI [0.01, 0.16]) (**Fig. 1e**), with invasive islands of neoplastic urothelial cells embedded in stroma and muscularis, exhibiting an irregular tumor-stroma interface and nuclear features progressing from low- to high-grade with depth. One sample (3.2%; 95% CI [0.01, 0.16]) displayed a tubular/microcystic subtype (**Fig. 1f**), characterized by small nests and tubules of neoplastic cells arranged in a puzzle piece-like pattern, with low-grade nuclear morphology and minimal stromal response despite extensive invasion.

Immunohistochemistry was performed on select cases and is summarized in **Table 2**. Antibodies included UPKIII (a marker of urothelial umbrella cells and typically luminal differentiation), E-cadherin (a transmembrane adhesion molecule), and vimentin (a mesenchymal marker and indicator of epithelial-mesenchymal transition).

**Table 2.**
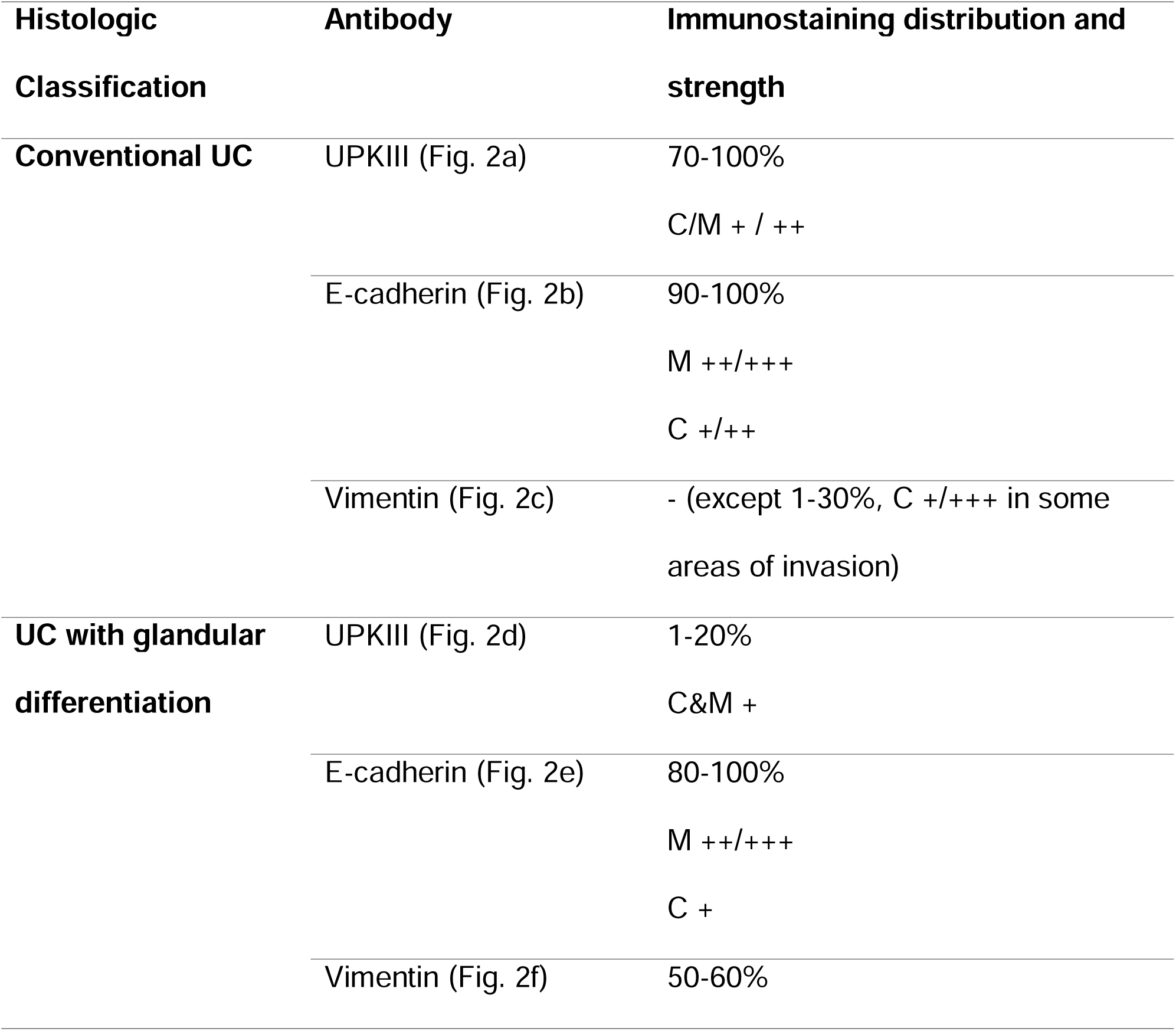
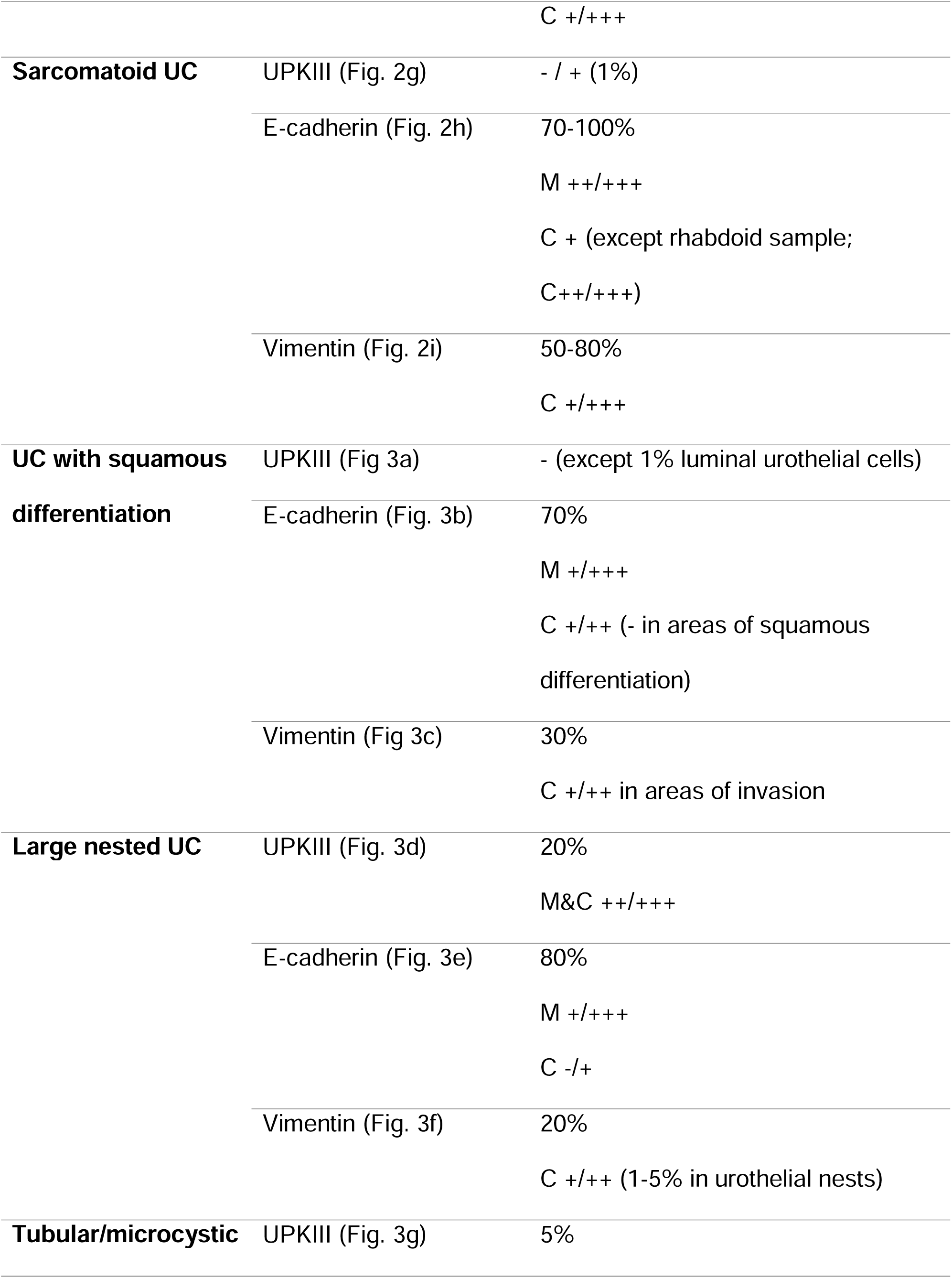
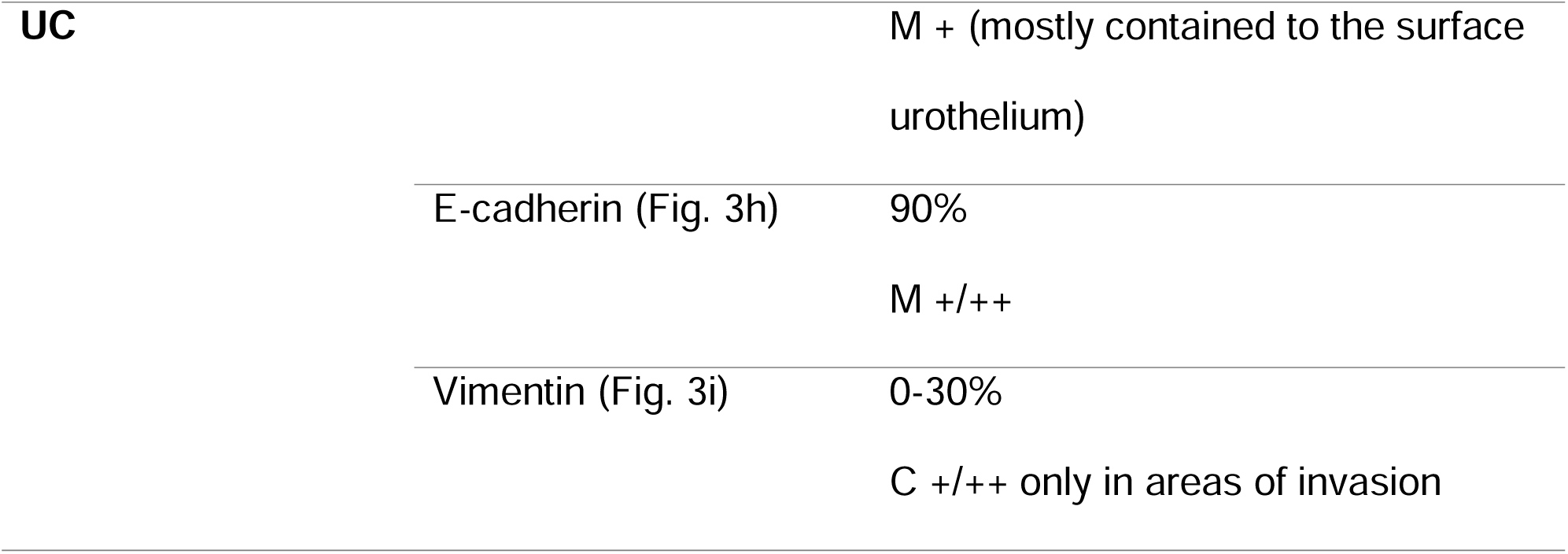
Summary of immunohistochemical (IHC) patterns of representative cases of canine urothelial carcinoma after classification of histologic subtypes and divergent differentiation. Distribution is described as cellular localization of staining (membranous (M), cytoplasmic (C), nuclear (N)) and pattern (rare (≤ 1%), sporadic (1-20%), patchy (20-79%), extensive (80-99%), diffuse (100%)), which was defined for this study as the estimated percentage of neoplastic cells in the section with positive immunostaining. Staining strength was subjective and graded as negative (-), faint (+), moderate (++), or strong (+++).

In conventional UC, UPKIII immunoreactivity was faint to moderate, patchy (70%) to diffuse, and predominantly cytoplasmic to membranous (**Fig. 2a**). E-cadherin expression was extensive (90%) to diffuse, with moderate to strong membranous and faint to moderate cytoplasmic staining (**Fig. 2b**). Notably, E-cadherin tended to be reduced in invasive areas or toward the lumen in papillary cases. Vimentin was largely absent in neoplastic cells but present in surrounding stroma (**Fig. 2c**); however, some tumors showed rare to patchy (1–30%) cytoplasmic expression, especially in large anaplastic cells at invasive edges.

**Figure 2.**
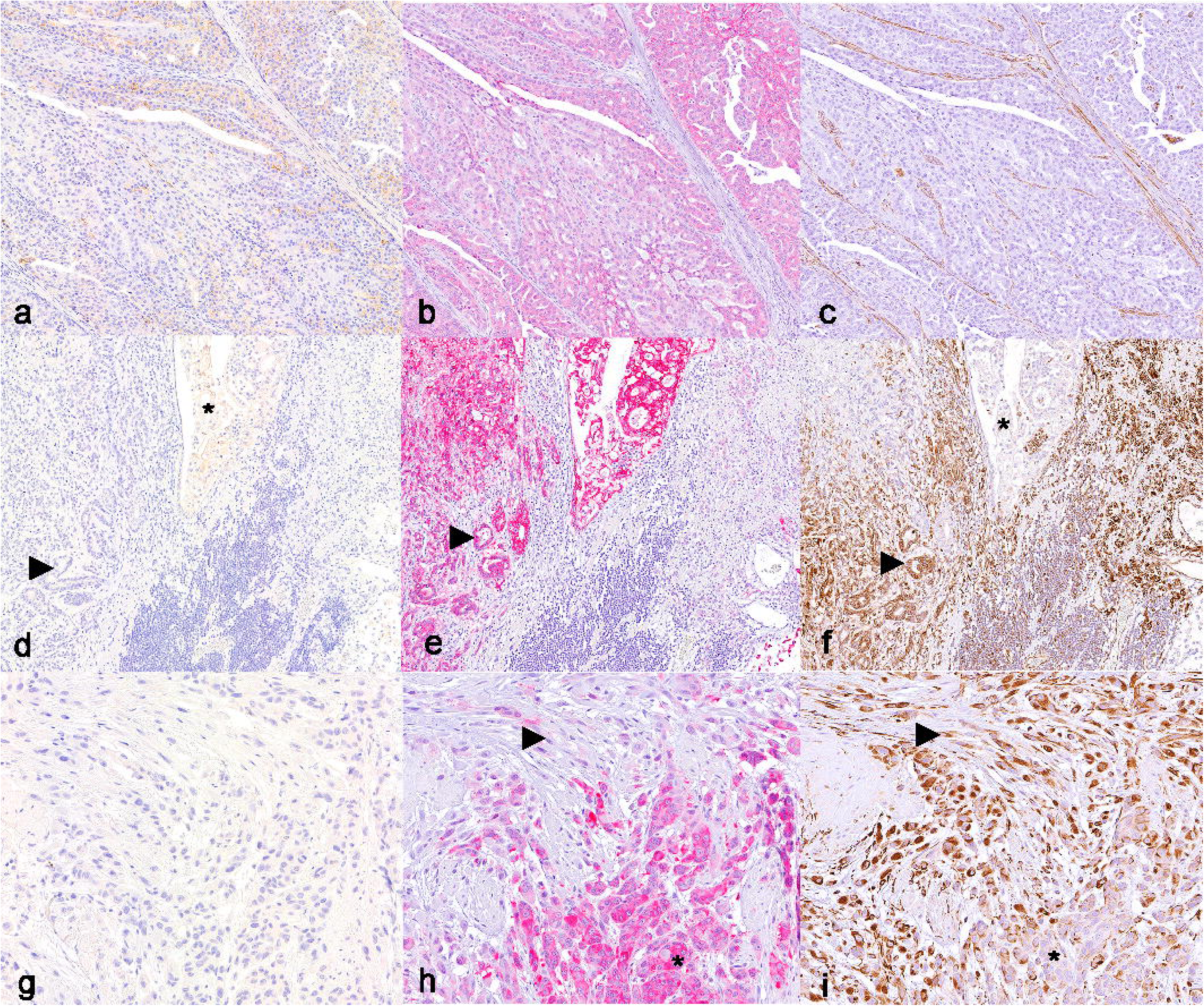
Immunohistochemistry of conventional UC and subtypes and divergent differentiations of UC identified in the urogenital tracts of dogs. Conventional UC, Case 7. a) Patchy areas of cytoplasmic uroplakin III immunoreactivity within neoplastic urothelial cells. b) Faint to moderate cytoplasmic E-cadherin immunoreactivity, with stronger membranous immunoreactivity in neoplastic cells toward the lumen. c) Neoplastic urothelial cells are diffusely immunonegative for vimentin. **UC with glandular differentiation, Case11.** d) Neoplastic urothelial cells forming more conventional islands have faint to moderate cytoplasmic uroplakin III immunoreactivity (asterisk), while those forming glands are immunonegative (arrowhead). e) Neoplastic cells have strong cytoplasmic E-cadherin immunoreactivity. f) Neoplastic urothelial cells forming more conventional islands are immunonegative for vimentin (asterisk), while those forming glands are immunopositive (arrowhead). **Sarcomatoid UC, Case 14.** g) Neoplastic urothelial cells are diffusely immunonegative for uroplakin III. h) E-cadherin immunoreactivity is moderate to strong and predominantly cytoplasmic in sheets of neoplastic urothelial cells (asterisk), with decreased to absent immunoreactivity in spindle-shaped urothelial cells invading deeper into the stroma (arrowhead). i) Vimentin immunoreactivity (a marker of epithelial-mesenchymal transition) is moderate to strong in spindle-shaped urothelial cells invading deeper into the stroma (asterisk), with decreased to absent immunoreactivity in more cohesive islands of neoplastic urothelial cells (arrowhead).

Tumors with extensive glandular differentiation located in the prostate underwent UPKIII staining to confirm urothelial origin. These showed faint, patchy to sporadic (1–20%) cytoplasmic and membranous staining, which was lost in gland-like structures (**Fig. 2d**). E-cadherin expression was extensive to diffuse, with moderate to strong membranous and faint cytoplasmic staining; in invasive regions, membranous staining diminished while cytoplasmic staining increased (**Fig. 2e**). Vimentin staining was patchy (50–60%) and ranged from faint to strong cytoplasmic (**Fig. 2f**).

Sarcomatoid UC had rare (1%) or absent UPKIII staining (**Fig. 2g**), with moderate to strong membranous staining only at the surface urothelium. E-cadherin was patchy (70%) to diffuse, with moderate to strong membranous and faint cytoplasmic expression, except in the rhabdoid case, which exhibited stronger cytoplasmic reactivity (**Fig. 2h**). E-cadherin expression declined in invasive areas. Vimentin expression was cytoplasmic, patchy (50%) to extensive, and ranged from faint to strong (**Fig. 2i**).

In UC with squamous differentiation, UPKIII was largely negative, with only rare (1%) faint cytoplasmic staining in luminal cells (**Fig. 3a**). E-cadherin expression was patchy (70%) with variable membranous and faint to moderate cytoplasmic expression, but diminished in squamous areas (**Fig. 3b**). Vimentin was patchy (30%), with faint to moderate cytoplasmic expression in invasive regions (**Fig. 3c**).

**Figure 3.**
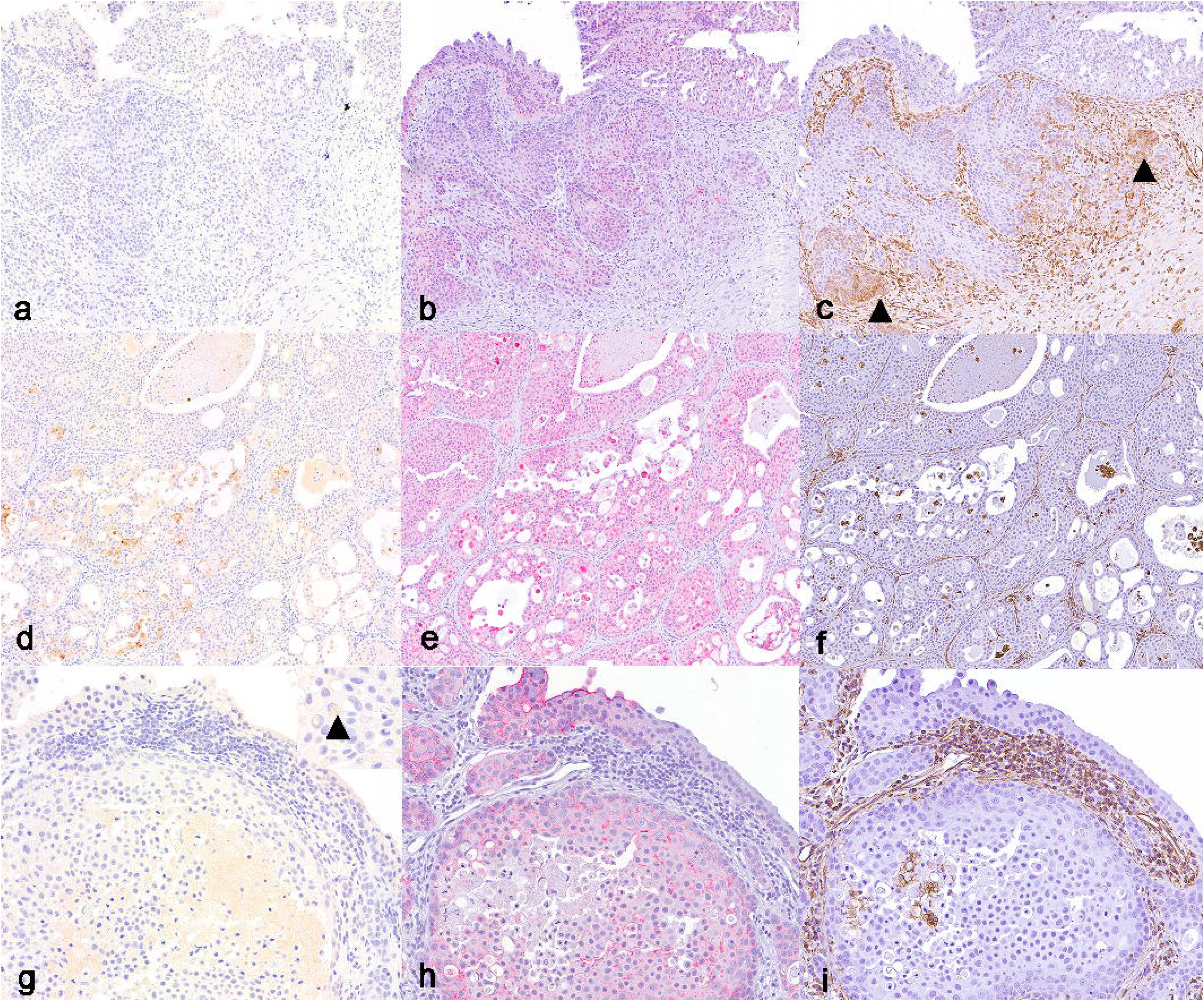
Immunohistochemistry of subtypes and divergent differentiations of UC identified in the urogenital tracts of dogs. UC with squamous differentiation, Case 28. a) Neoplastic urothelial cells are predominantly immunonegative for uroplakin III. b) Faint to moderate cytoplasmic E-cadherin immunoreactivity, with decreased expression in areas of squamous differentiation. c) Most neoplastic urothelial cells are vimentin immunonegative, with cytoplasmic expression only in areas of invasion (arrowheads), suggestive of epithelial to mesenchymal transition. **Large nested UC, Case 30.** d) Large nests of neoplastic urothelial cells within the stroma have patchy areas of moderate to strong membranous and cytoplasmic immunoreactivity. e) E-cadherin immunoreactivity within the neoplastic nests is faint to strong, membranous and negative to faintly cytoplasmic. Rare cells have strong cytoplasmic expression. f) Vimentin immunoreactivity within the neoplastic nests is predominantly negative, only highlighting a few dissociated urothelial cells that have sloughed into the cystic lumens. **Microcystic/tubular UC, Case 31.** g) Neoplastic urothelial cells are immunonegative for uroplakin III, with rare immunoreactivity within Melamed-Wolinska bodies (inset, arrowhead). h) E-cadherin immunoreactivity is extensive, with faint to moderate membranous expression. i) Vimentin immunoreactivity is predominantly negative.

Large nested UC had patchy (20%), moderate to strong membranous and cytoplasmic UPKIII staining (**Fig. 3d**). E-cadherin was extensive (80%), with variably faint to strong membranous expression and predominantly negative to faint cytoplasmic expression (**Fig. 3e**). Vimentin staining was patchy (20%), with faint to moderate cytoplasmic staining that increased in invasive areas (**Fig. 3f**).

Tubular/microcystic UC had sporadic (5%), faint membranous UPKIII staining, mostly confined to the surface urothelium (Fig. 3g), with occasional Melamed-Wolinska bodies (**Fig. 3g**, inset), but was otherwise immunonegative. E-cadherin was extensive (90%), with faint to moderate membranous staining strongest at the surface (**Fig. 3h**). Vimentin was largely negative, with patchy (30%) faint to moderate cytoplasmic staining in invasive regions (**Fig. 3i**).

Data on the presence or absence of metastatic disease, when available, are summarized in **Fig. 4**. Ten cases did not have sufficient information in the clinical record regarding metastasis and were excluded from analysis. Notably, cases of UC with glandular differentiation had a higher incidence of metastasis compared to conventional UC. Survival data for all UC cases are presented in **Fig. 5**, both as a Kaplan–Meier curve (top) and stratified by UC subtype or divergent differentiation(bottom). Six cases did not have sufficient information in the clinical records and were excluded from analysis. The median survival time across all cases, ignoring any potential effect of subtype or divergent differentiation, was 9 months (95% CI: 6–∞).

**Figure 4.**
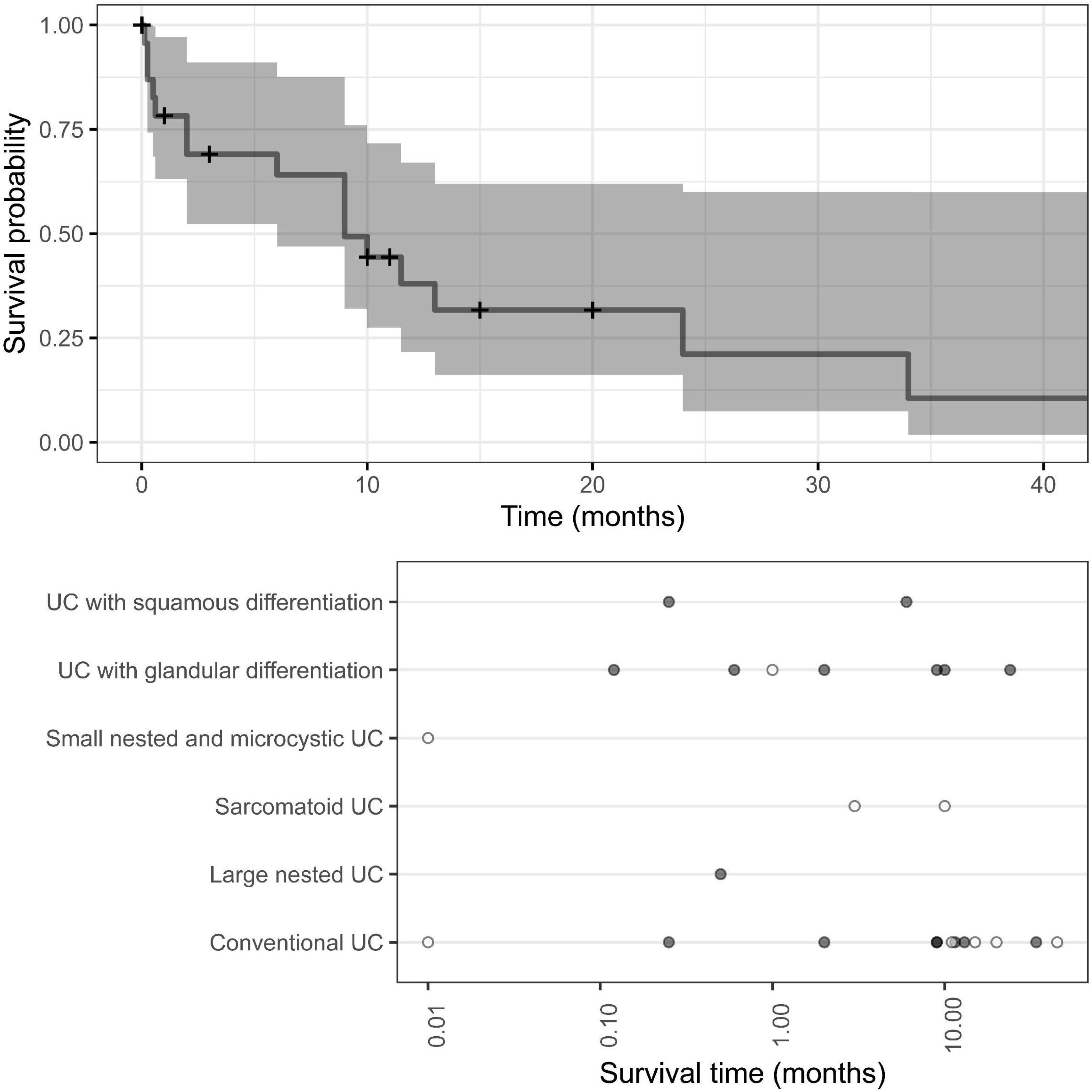
Apparent metastasis prevalence in canine UC. Count (upper panel) and proportion (lower panel) of metastasis-positive and negative cases by histologic subtype and divergent differentiation. Pale gray indicates metastasis-positive as determined by radiographs or necropsy, while dark gray indicates metastasis-negative. Error bars indicate posterior median predicted count and 90% credible interval, determined from a hierarchical logistic model implementing between-histologic type variation.

**Figure 5:**
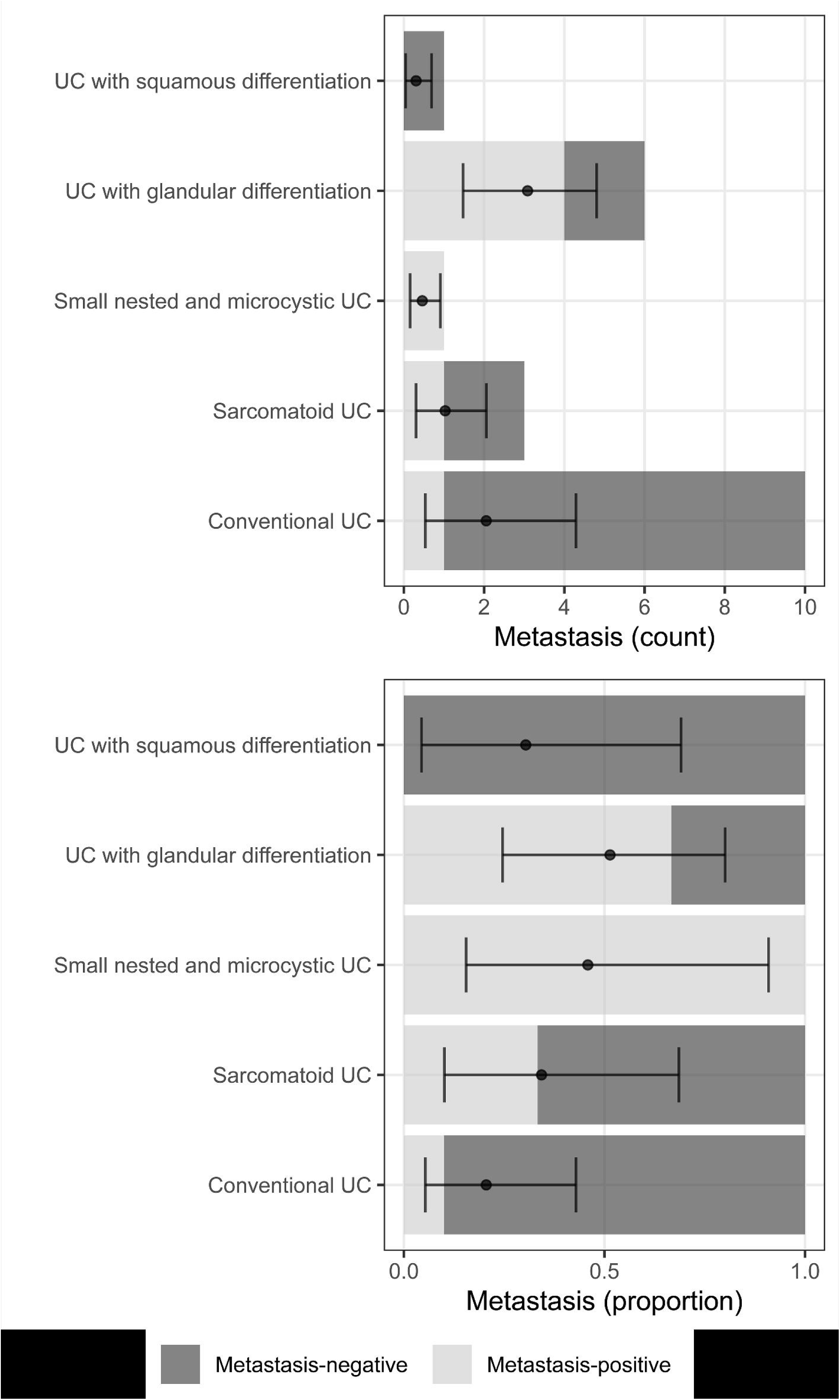
Survival time after presentation for 31 UC cases. The upper panel describes the observed survival time, across all subtypes. The solid line is the Kaplan-Meier estimate of the survival function, and the gray field is its 95% confidence region. Note that the upper 95% confidence limit of the median survival time is ∞. The lower panel describes the observed survival time by subtype, where the filled points are the survival time for cases observed until death, and the open circles are cases that were lost-to- follow-up (censored).

## DISCUSSION

Recent efforts by the WHO to incorporate histological subtypes into the classification system of human UC have underscored the importance of routinely identifying and reporting these variants. Many of the UC subtypes, particularly sarcomatoid, are associated with aggressive clinical behavior, frequent and early metastases, poor responses to chemotherapy, and reduced overall survival ^7^. In this study, the histology of canine UC cases from multiple institutions was reviewed, focusing on the subtypes and divergent differentiation with the expertise of a human uropathologist. This approach resulted in the reporting of a higher number of sarcomatoid UC (12.9%) and UC with glandular differentiation (25.8%) than previous reports in both human and canine cohorts (sarcomatoid UC: Human 0.3-7%, canine: 3.4%; UC with glandular differentiation: Human 1.8-2.6%, canine: 2%; see **Table 1**). This increased detection may reflect the added diagnostic precision afforded by combined pathology expertise, despite a smaller sample size compared to previous canine studies (n=260 ^14^).

However, percentage or incidence reports are not always directly comparable between studies, as they heavily depend on the type of cases included in the study population, as well as the overall number of cases. For example, samples obtained through trans-urethral resection of bladder tumors (TURBT) in humans are more likely to be performed on lower grade and stage conventional bladder cancer than those obtained by radical cystectomy ^1,63^. In addition, discordance between TURBT and cystectomy histology are well-documented with TURBT showing conventional UC while the cystectomy contains subtype component ^1,55,63^. Smaller, fragmented, or more superficial samples obtained via TURBT may limit the accurate identification of subtypes as well ^32^. A similar limitation may apply to canine cases; cystotomy or endoscopic samples are often superficial and fragmented, whereas necropsy specimens provide larger, more representative sections of the tumor. Additionally, certain histologic subtypes, including sarcomatoid and plasmacytoid variants, are thought to result from late-stage dedifferentiation of basal or luminal UC ^30,60^, suggesting that tumor “biological age” plays a role in subtype emergence. Therefore, unless the study population is stratified by invasiveness or tumor stage (e.g., MIBC vs. non-MIBC), reported subtype frequencies may be skewed. Given that dogs have a higher prevalence of muscle-invasive bladder cancer than humans (approximately 60,000 vs. 16,000 cases per year)^4,16^, the canine population offers a valuable opportunity for comparative studies, particularly of rare and aggressive subtypes. Our cohort may also have been enriched for more advanced disease, as necropsy specimens comprised ∼40% of the cases (**Supplemental Table S1**). By contrast, a prior large-scale canine study focused on diagnostic samples could have had more superficial tumor samples, akin to TURBT samples in human patients (**Supplemental Table S1**). Additional factors that have been implicated in driving the development of histological subtypes in humans are treatment with certain chemotherapies, such as cisplatin, which is a commonly used first-line treatment in human MIBC ^6^.

Due to the low number of each subtype in this study, evaluation of correlation between treatment, outcomes, and the occurrence of histological subtypes in the case series could not be performed. It would, however, be interesting to follow the natural progression of the disease in canine breeds that are at a higher risk of developing bladder cancer, and/or to interpret multiple biopsies from the same patients before and after chemotherapy in the future. The overall number of samples included in the study population is also important, as seen in some very large studies using the NCI’s Surveillance, Epidemiology, and End Results Program (SEER) database on human cancers ^65^, which are likely much more informative in regards to the true incidence of these histological subtypes than the smaller studies reported in veterinary medicine to date.

With regard to specific histological subtypes, we report positive evidence of a higher proportion of sarcomatoid UC subtypes (12.9%) than currently published data in humans and dogs. This subtype is associated with poor survival in humans ^17^. In our study, there was an insufficient number of cases to determine if there was greater incidence of metastasis in cases identified as having a sarcomatoid UC subtype. Efforts to molecularly subtype these tumors in humans have identified that the sarcomatoid regions of the tumor are likely derived from basal UC stem cells expressing markers such as CD44, CK5/6, and CK14 ^19^. In addition, tumor regions that are histologically identified as sarcomatoid also express increasing amounts of vimentin and N-cadherin, while having decreased expression of E-cadherin. This suggests a transition of UC basal stem cells to a less adherent and more invasive phenotype. The latter expression signature is a hallmark of epithelial-to-mesenchymal transition (EMT) ^19^ and, critically, has been associated with increased metastatic potential and aggressiveness in other tumors having high heterogeneity ^26^. Similarly, we show here that regions of tumor identified as a sarcomatoid subtype of canine UC were highly positive for vimentin, with both aberrant (cytoplasmic) and decreased E-cadherin expression (**Fig. 2h, i**).

Furthermore, there was evidence of a higher frequency of UC with glandular differentiation in our study (25.8%) than previously reported in humans (1.8-2.6%) and in dogs (2%). A glandular component of urothelial carcinoma is often characterized by the formation of glandular structures or it can recapitulate the appearance of enteric histology, resembling a colonic-type adenocarcinoma, or, alternatively, the presence of mucinous type carcinoma with mucin pools containing either glands or signet ring cells ^42^. The presence of glandular morphology results in the acquisition of an alternate immuno-phenotype with expression of CK20 and CDX-2, typical of enteric lesions with either co-expression or loss of urothelial markers ^54^. In this study, we identified expression of uroplakins, which confirms the cellular identity of the tumor as UC with glandular differentiation as opposed to adenocarcinoma, with loss of UPKIII expression in areas with glandular arrangement (**Fig. 2d**).

In the present cohort, 6% of cases were UC with squamous differentiation, which is similar to what has been reported in humans (8-13%), and canines (3%) (**Table 1**). Histologically, squamous differentiation is defined by the presence of keratinization and intercellular bridges, often seen in the setting of chronic irritation from bladder stones ^11^, *Schistosoma* infection (humans) ^56^, or neurogenic bladder with indwelling catheters ^21^. As for markers, human UC with squamous differentiation often have positive expression for basal urothelial cell markers such as CK5/6, CK14, and p63, and negative expression for CK20 and uroplakins ^18^. In this study, only one sample was able to be analyzed due to tissue size and was predominantly negative for uroplakin III with the exception of rare luminal urothelial cells, which implies loss of luminal urothelial differentiation (**Fig. 3a**).

In this study, there was one case each of microcystic/tubular and large nested histological subtypes of UC. These cases appeared to have a similar prevalence in our dog population as previously reported in humans, although we only observed one case of each subtype. In humans, the nested and microcystic/tubular histological subtypes are rare but important to recognize, as they have deceptively benign histology ^58^. The tubular/microcystic subtype features prominent microcysts, macrocysts, or tubular structures with lumens that are usually lined by simple cuboidal epithelium ^37^. There are conflicting reports on the lineage derivation of this subtype, with recent cases showing both basal and luminal markers ^58^. The prognosis in human cases is generally poor, although it may be similar to the reported survival in conventional UC cases ^58^.

The case signalment in this current study was very similar to what has previously been described for canine UC ^14,16^, with a median age of 11 years (range: 7-15 years). Limitations of the study include the relatively small sample size, limited archival material in some cases, and the inability to correlate histological findings with clinical data. While we observed a higher frequency of metastasis in UC with glandular differentiation, the sample size was insufficient for definitive interpretation. Similarly, although the median survival time for UC cases was 9 months, we could not determine whether survival differed by histologic subtype.

In conclusion, we identified histological subtypes in 31 cases of canine bladder cancer and found higher-than-expected frequencies of sarcomatoid UC and UC with glandular differentiation. These findings support the inclusion of histological subtype reporting in canine UC, following human diagnostic frameworks. Moreover, the overrepresentation of sarcomatoid UC and UC with glandular differentiation in dogs suggests that canine models may offer a valuable, naturally occurring platform for studying rare UC subtypes and testing novel therapeutic strategies, especially for aggressive variants like sarcomatoid UC and UC with glandular differentiation ^43,72^ that are exceedingly rare in humans.

## Acknowledgments

We appreciate the timely processing of samples at the Histology and Pathology Departments at Iowa State University, University of Georgia, Cornell University, and Purdue University. The authors acknowledge Dr. Deborah Knapp in the Werling Comparative Oncology Research Center at Purdue University for contributing case samples and information. The authors also acknowledge the assistance of Megan Cohen and the Purdue University Histology Research Laboratory, a core facility of the NIH-funded Indiana Clinical and Translational Science Institute.

## CRediT authorship contribution statement

Megan P. Corbett: Resources acquisition, Investigation, Data curation, Formal analysis,

Writing – review & editing

Mohamed Elbadawy: Investigation, Methodology, Data curation, Formal analysis,

Writing – original draft, Writing – review & editing

Karin Allenspach: Conceptualization, Funding acquisition, Supervision, Project administration, Methodology, Investigation, Data curation, Formal analysis, Writing – original draft, Writing – review & editing

Hannah Wickham: Methodology, Resources acquisition, Investigation, Data curation,

Formal analysis, Writing – review & editing

Hannah Nicholson: Methodology, Resources acquisition, Writing – review & editing

Michael Catucci: Methodology, Resources acquisition, Writing – review & editing

Bryan J. Melvin: Methodology, Resources acquisition, Writing – review & editing

Basant Ahmed: Methodology, Resources acquisition, Investigation, Data curation, Formal analysis, Writing – review & editing

Christopher Zdyrski: Methodology, Resources acquisition, Investigation, Data curation, Formal analysis, Writing – review & editing

Lilian J. Oliveira: Resources acquisition, Investigation, Data curation, Formal analysis Writing – review & editing

Elizabeth W. Howerth: Resources acquisition, Investigation, Data curation, Formal analysis Writing – review & editing

John, C. Cheville: Conceptualization, Supervision, Methodology, Investigation, Writing – review & editing

Aleksandra Pawlak: Methodology, Writing – review & editing

Olufemi Fasina: Resources acquisition, Investigation, Data curation, Formal analysis Writing – review & editing

Jonathan P. Mochel: Funding Acquisition, Supervision, Project administration, Methodology, Writing – review & editing.

Andrew Woodward: Methodology, Formal analysis, Writing – review & editing

## Supplementary Material

Please find the following supplemental material available below.

Supplemental Table S1

Supplemental Table S2

