## Supplemental Table 1 for "Frequency and Characterization of Rare Histologic Subtypes in Canine Invasive Urothelial Carcinoma"

**Supplemental Table S1.** Canine patient information.

| **Case** | **Source** | **Age (Y)** | **Breed** | **Sex** | **Necropsy/**  **Biopsy** | **Sample site** | **Histological Review** | **Metastasis** |
| --- | --- | --- | --- | --- | --- | --- | --- | --- |
| 1 | Iowa State University | 14 | Mixed breed | mn | N | Urinary bladder | Conventional UC | N/A |
| 2 |  | 14 | Mixed breed | fs | N | Urinary bladder/proximal urethra | Conventional UC | No |
| 3 |  | 13 | Maltese | mn | N | Urinary bladder | Conventional UC | No |
| 4 |  | 11 | Mixed breed | mn | N | Prostatic urethra | Conventional UC | No |
| 5 |  | 9 | Bernese mountain dog | fs | N | Urethra | Conventional UC | Yes |
| 6 |  | 12 | Yorkshire terrier | fs | N | Urinary bladder | Conventional UC (papillary) | No |
| 7 |  | 11 | Shepherd mix | mn | B | Urinary bladder | Conventional UC (papillary) | N/A |
| 8 |  | 11 | Mixed breed | mn | N | Urinary bladder | Conventional UC | No |
|  |  |  |  |  |  | Urinary bladder | UC with glandular differentiation |  |
| 9 |  | 9 | Mixed breed | mn | N | Prostatic urethra/prostate | UC with glandular differentiation | Yes |
| 10 |  | 15 | Border Collie mix | mn | N | Prostatic urethra | UC with glandular differentiation | Yes |
| 11 |  | 12 | German Shepherd dog | mn | N | Prostate gland | UC with glandular differentiation | Yes |
| 12 |  | 15 | Labrador retriever | m | N | Renal pelvis | UC with squamous differentiation and sarcomatoid features | No |
| 13 |  | 12 | Labrador retriever | fs | N | Urethra | Sarcomatoid UC | No |
| 14 |  | 12 | N/A | f | B | Urinary bladder | Sarcomatoid UC w/ rhabdoid features | N/A |
| 15 | Purdue University | 11 | Mixed breed | fs | B | Urethra | Conventional UC (papillary) | No |
|  |  | 12 |  |  | B | Urethra | Conventional UC (papillary) |  |
| 16 |  | 13 | Pomeranian | f | B | Urinary bladder | UC in situ with glandular differentiation | No |
| 17 |  | 11 | Mixed breed | fs | B | Urethra/urinary bladder | Sarcomatoid UC | No |
| 18 | University of Georgia | 9 | Labrador Retriever | fs | B | Urinary bladder | Conventional UC | N/A |
| 19 |  | 7 | Coonhound Mixed breed | fs | B | Urinary bladder | Conventional UC | No |
| 20 |  | 11 | Beagle mix | mn | B | Urinary bladder | Conventional UC (papillary) | No |
| 21 |  | 8 | Brittany Spaniel | fs | B | Urinary bladder | Conventional UC (papillary) | No |
| 22 |  | 12 | Pomeranian | fs | B | Urinary bladder | Conventional UC (papillary) | No |
| 23 |  | 11 | Swiss Mountain Dog | fs | B | Urinary bladder | Conventional UC (papillary) | N/A |
| 24 |  | 12 | Mixed breed | mn | B | Urinary bladder | Conventional UC, exophytic and inverted growth pattern (papillary) | N/A |
| 25 |  | 11 | Maltese | mn | B | Urinary bladder | UC with glandular differentiation | N/A |
| 26 |  | 10 | Havanese | mn | B | Urinary bladder | UC with glandular differentiation (papillary) | N/A |
| 27 |  | 9 | English Springer Spaniel | fs | B | Urinary bladder | UC with glandular differentiation and exophytic growth pattern (papillary) | Yes |
| 28 |  | 10 | English bulldog | fs | B | Ureter | UC with squamous differentiation | Yes |
| 29 |  | 11 | Dachshund | fs | B | Urinary bladder | Sarcomatoid UC | Yes |
| 30 |  | 10 | Cavalier King Charles Spaniel | mn | B | Prostatic metastasis of primary tumor in bladder | Large nested UC | N/A |
| 31 |  | 10 | Alaskan Malamute | fs | B | Urinary bladder | Small nested and microcystic UC with squamous metaplasia | Yes |

UC: urothelial carcinoma, m: intact male, f: intact female, mn: male neutered, fs: female spayed, N: necropsy, B; biopsy, N/A: not applicable as no data is available.
