## Supplemental Table 2 for "Frequency and Characterization of Rare Histologic Subtypes in Canine Invasive Urothelial Carcinoma"

**Supplemental Table S2**. Immunohistochemistry markers for further characterization of 31 cases of canine urothelial carcinoma.

| Antibody | Marker | Host | Source, Catalog Number | Clone | Antigen Retrieval | Dilution | | Positive Control |
| --- | --- | --- | --- | --- | --- | --- | --- | --- |
| Uroplakin III | Urothelial luminal (umbrella) cells | Mouse | Fitzgerald Industries  10R-U103ax | AU1 | Bond epitope retrieval solution 2 (Leica) for 40 min at RT | | 1:20 for 10 min at RT | Urinary bladder |
| E-cadherin | Cellular adhesion and polarity | Mouse | BD Biosciences 610181 | 36/E-Cadherin | Citrate (pH 6.0) for 15 min at 110°C | | 1:500 for 60 min at RT | Haired skin |
| Vimentin | Mesenchymal origin (epithelial - mesenchymal transition) | Mouse | BioGenex MU074-UC | V9 | Citrate (pH 6.0) for 15 min at 110°C | | 1:3000 for 60 min at RT | Small intestine |

RT = room temperature
